# Semantic representations in the visual cortex of blind and sighted humans

**DOI:** 10.1101/2025.01.28.635293

**Authors:** Małgorzata Paczyńska, Marta Urbaniak, Marta Dębecka, Łukasz Bola

## Abstract

In blind humans, the “visual” cortex responds to linguistic stimuli, such as words and sentences. This is sometimes taken as evidence that this brain region supports starkly different computations in blind and sighted individuals. Here, we challenge this view and show that, during word processing, the visual areas in these two populations represent the same semantic dimension – the knowledge about physical properties of word referents. Using analysis of fMRI activation patterns, we found that the visual cortex in both congenitally blind and sighted participants represented differences between individual words. In both groups, the activation patterns for words in the visual cortex reflected physical, but not conceptual similarity between word referents. Furthermore, the between-group correlations in these activation patterns were comparable to within-group correlations. Finally, during word processing, the visual areas in both groups showed greatest “representational connectivity” to the occipitotemporal areas. Overall, our findings suggest that responses to linguistic stimuli in the visual cortex of blind individuals are driven by representational mechanisms that are functional also in the sighted adult brain. In sighted individuals, information about physical properties of word referents might be backprojected to visual areas, from the occipitotemporal cortex, to support visual predictions, imagery, and visuospatial thinking. In blind individuals, this mechanism might be preserved and, combined with increased excitability of the blind visual cortex, drive strong responses of this region to linguistic stimuli.

## Introduction

Language processing involves comparable brain regions across languages and cultures (Malik-Moraleda et al., 2022). These are primarily areas clustered around the left sylvian fissure, such as the inferior frontal gyrus, the superior temporal lobe, and the anterior temporal lobe (Fedorenko et al., 2024).

Intriguingly, in blind individuals, language processing activates not only the canonical language network, but also the “visual” cortex. (Sadato et al., 1996; Burton et al., 2002; Röder et al., 2002; Bedny et al., 2011; Bedny et al., 2015; Lane et al., 2015). The magnitude of these activations in the visual areas of blind individuals increases with increasing complexity of linguistic stimuli (Röder et al., 2002; Bedny et al., 2011; Lane et al., 2015), and is greater for semantic tasks than for phonological tasks (Burton et al., 2003). Transient disruption of the visual cortex activity in blind individuals interferes with certain operations involving linguistic stimuli, such as Braille reading (Cohen et al., 1997, Kupers et al., 2007) or word generation (Amedi et al., 2004).

An increasingly popular interpretation of these findings is that they signify a profound change in the implementation of cognitive functions in the brain, induced by the absence of visual experience (Bedny, 2017; Saccone et al., 2024). As a result of this process the visual cortex in blind humans might potentially support high-level cognitive functions, which are not supported by visual areas of the sighted.

However, an alternative possibility is that responses to linguistic stimuli in the blind visual cortex are driven by neural mechanisms that are functional also in the sighted adult brain. The sighted visual cortex receives information from the higher-level brain regions (Roelfsema and de Lange, 2016). Thus, signals from the language network can presumably modulate the activity of the visual cortex also in sighted individuals. Certain visual areas in the sighted are activated by spoken words and sentences (Seydell-Greenwald et al., 2023) and represent physical properties of word referents (Borghesani et al., 2016; Martin et al., 2018). At least some of these effects are unlikely to be driven by visual imagery (Seydell-Greenwald et al., 2023). Based on these results one may suppose that, following language comprehension, the knowledge about physical properties of word referents is backprojected to the visual system in an attempt to predict incoming visual signals (Rao and Ballard, 1999; Lee and Mumford. 2003; Friston, 2005). In this study, we hypothesized that this mechanism develops even in blind individuals and, in the absence of visual inputs, drives strong responses of the blind visual cortex to linguistic stimuli. The comparable representations of physical properties of word referents has been already documented in the occipitotemporal areas of blind and sighted individuals (Mahon et al., 2009; He et al., 2013; Peleen et al., 2013, 2014; Xu et al., 2023). Here, we asked whether such representations can drive responses to linguistic stimuli in the occipital lobe, particularly in the early visual areas, which has been suggested to be a hotspot of functional plasticity in the blind brain (Amedi et al., 2003).

To investigate this issue, we performed an fMRI study in which sighted and congenitally blind individuals were presented with 20 concrete words. The words were presented in either the spoken or the written modality (the visual alphabet in the sighted group and the Braille alphabet in the blind group), and the participants were asked to answer questions about either physical (e.g., is this round?) or conceptual (e.g., is this living?) properties of word referents (Martin et al., 2018; Xu et al., 2023). Furthermore, in a separate behavioral study, we collected the ratings of physical and conceptual similarity between word referents for all words used in the fMRI study.

We used the analysis of fMRI activation patterns to investigate neural representations induced in the visual areas of blind and sighted participants. If responses to linguistic stimuli in the blind visual cortex are driven by a change in cognitive functions implemented in this region, then the visual areas in both groups should represent different word properties. One can particularly expect that the visual cortex in the blind, but not in the sighted individuals, represents abstract, conceptual aspects of word meaning. Conversely, if the responses to linguistic stimuli in the blind visual cortex are driven by mechanisms that are present also in the sighted adult brain, one should find the same type of representations in the visual areas in both groups. Based on our hypothesis, this should be primarily the representation of the physical properties of word referents, which, in sighted individuals, can be particularly useful for visual predictions.

## Results

### Activation patterns in the visual cortex

We first asked whether the visual areas in blind and sighted individuals represent differences between specific concrete words. To address this question, we tried to decode individual words used in the study from the activation patterns of three occipital Brodmann areas (BAs): area 17 (primary visual cortex), area 18 (secondary visual cortex), and area 19 (associative visual cortex) (Figs. 1-2). To attenuate the impact of word sensory properties on the results, the decoding was performed across the modalities of word presentation, that is, the algorithm was always trained on one sensory modality and tested on the other. Generalization across the presentation modalities is one of the hallmarks of semantic representations in the higher-level brain regions (Liuzzi et al., 2017; Deniz et al., 2019).

**Figure 1.**
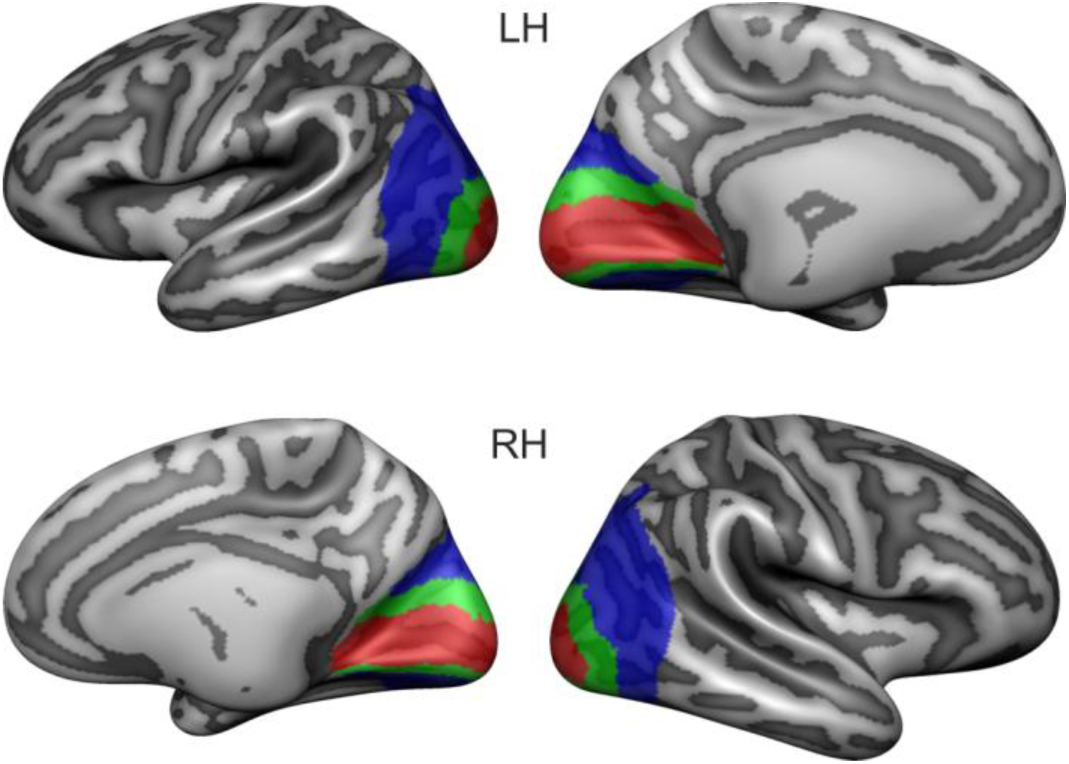
The masks of visual areas used in the study. The masks of Brodmann areas 17 (primary visual cortex, red), 18 (secondary visual cortex, green), and 19 (associative visual cortex, blue) are presented on the inflated reconstruction of participants’ averaged cortical anatomy.

**Figure 2.**
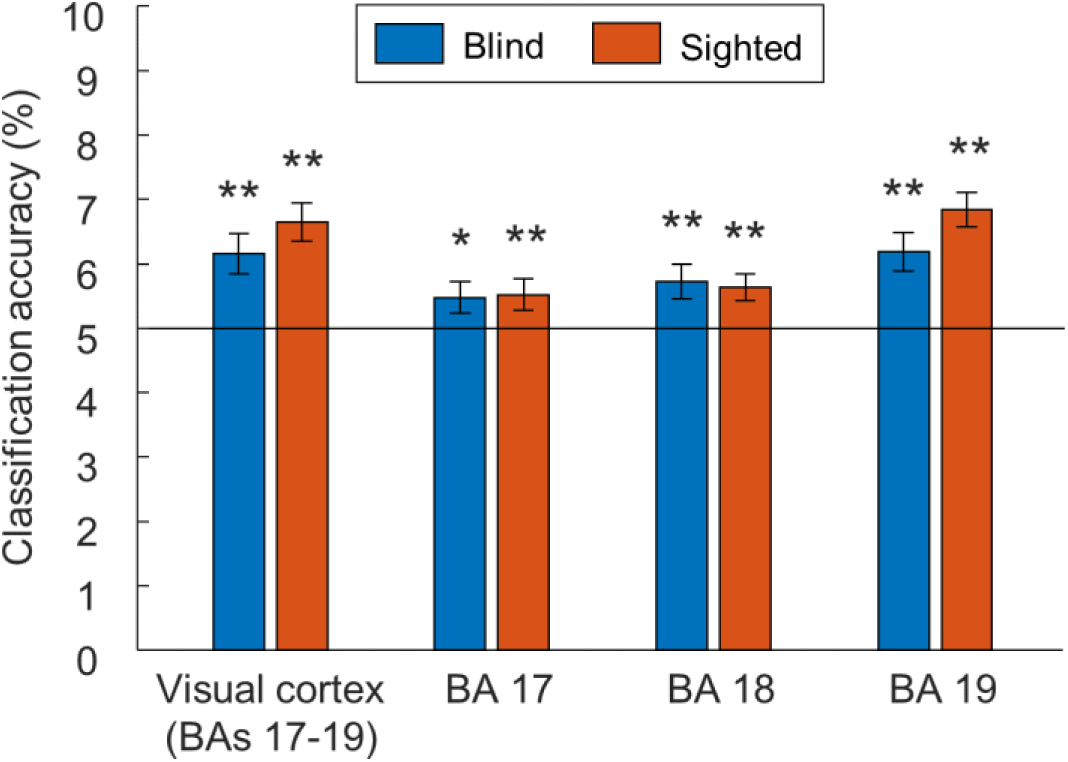
The visual cortex in blind and sighted individuals represents differences between individual concrete words. The results of cross-modal classification of activation patterns for individual words, presented in the spoken and the written modality, in the visual areas of blind and sighted participants. * p < 0.05, ** p < 0.01, corrected for multiple comparisons using the false discovery rate. Error bars represent the standard error of the mean. The black line indicates the chance classification level.

In both blind and sighted participants, the accuracy of crossmodal word decoding was significantly above chance level in the visual cortex (BAs 17, 18, and 19 combined; both p values < 0.01) and in all specific visual areas (all p values < 0.05). The 2 (group) x 3 (visual area) ANOVA produced only a significant main effect of the visual area (F(2,90) = 13.21, p < 0.001, partial eta squared = 0.23), with post-hoc tests indicating that the word decoding was more accurate in BA 19 than in the other two visual areas (both p values < 0.01). Neither the main effect of the group nor the interaction between the group and the visual area were significant (both p > 0.15). These results show that the visual cortex in both blind and sighted individuals represents differences between individual concrete words in a format that is independent of word presentation modality.

We then averaged the activity patterns for each word across the presentation modalities, and correlated similarity in these average activity patterns in the visual cortex with the behavioral ratings of physical and conceptual similarity between the word referents (Fig. 3; see Fig. S1 for the visualization of the behavioral similarity matrices). Since the two types of ratings were correlated (Spearman’s rho = 0.41, p < 0.001), we used a partial correlation approach, in which the results were calculated after accounting for this common variance.

**Figure 3.**
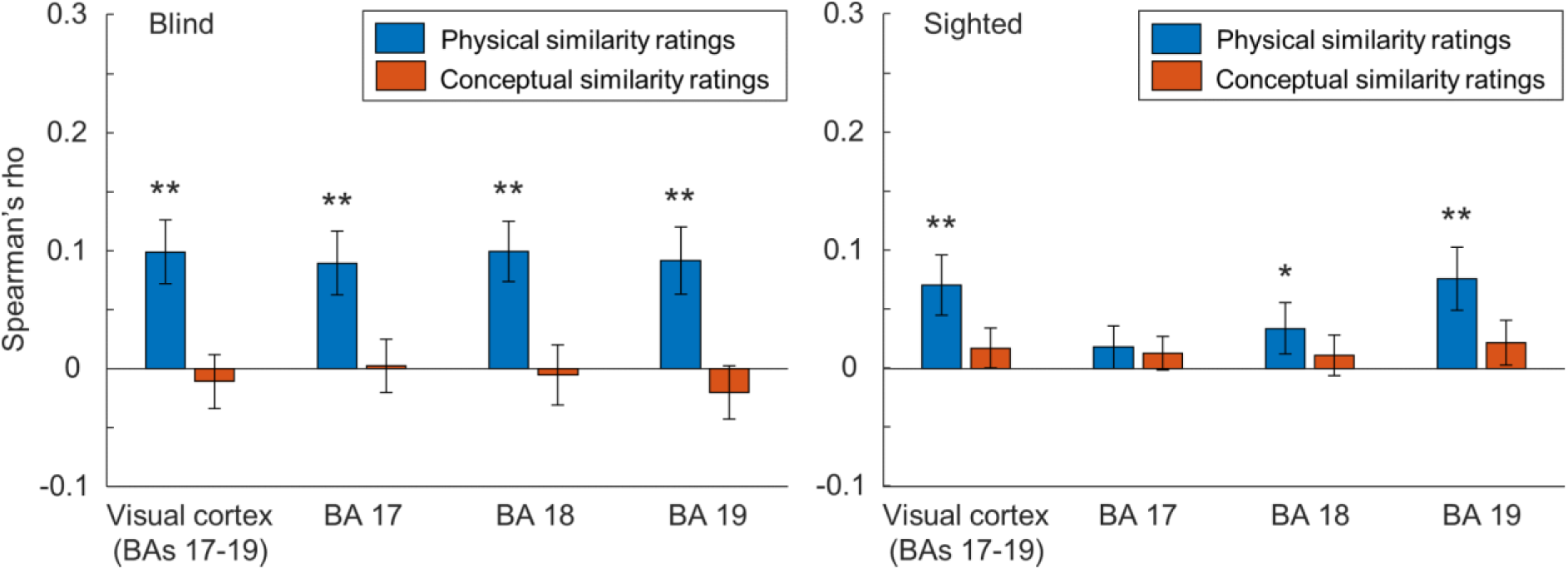
The activation patterns in the visual cortex of blind and sighted individuals reflect physical, but not conceptual similarity between word referents. Correlations between the neural similarity matrices, reflecting similarity between the activation patterns for individuals words in the visual areas of blind and sighted participants, and the behavioral ratings of physical and conceptual similarity between the word referents. * p < 0.05, ** p < 0.01, corrected for multiple comparisons using the false discovery rate. Error bars represent the standard error of the mean.

In both blind and sighted participants, the similarity in activity patterns for words in the visual cortex (BAs 17, 18, and 19 combined) was correlated with ratings of physical similarity of word referents (both p values < 0.01), but not with ratings of more abstract, conceptual similarity (both p values > 0.25). Significant correlations with ratings of physical similarity were found in all visual areas in the blind participants (all p values < 0.01) and in BAs 18 and 19 in the sighted participants (both p values < 0.05). In contrast, correlations with ratings of conceptual similarity were not observed in any visual area in either group (all p values > 0.25). In line with these results, the 2 (group) x 2 (similarity metric) x 3 (visual area) ANOVA produced a significant main effect of similarity metric (F(1,45) = 7.23, p = 0.01, partial eta squared = 0.14), which did not interact with the group or the visual area (both p values > 0.12). Additionally, the ANOVA also indicated a significant group by visual area interaction (F(2,90) = 6.79, p = 0.005, partial eta squared = 0.13), with post-hoc tests indicating that the correlation values were generally higher in BA 19 than in the other visual areas in the sighted participants (both p values < 0.01), but not in the blind participants (both p values > 0.25). No other main effects or interactions were significant. In summary, our results show that, in both blind and sighted individuals, the activity patterns for words in the visual cortex reflect the physical similarity of word referents, but not the more abstract, conceptual similarity.

While the three-way ANOVA interaction (group x similarity metric x visual area) on correlation scores was not significant (F < 1, p > 0.25), the activity patterns in BA 17 correlated with the ratings of physical similarity of word referents only in the blind group. We performed additional pairwise comparisons to further investigate this effect. The direct between-group comparisons confirmed greater correlation between the activity patterns in BA 17 and the ratings of physical similarity of word referents in the blind participants, compared to the sighted participants (t(45) = 2.28, p = 0.028, Cohen’s d = 0.67). No between-group differences were observed in other visual areas (both p > 0.05). The within-group comparisons showed that, in the sighted participants, the correlation with the ratings of physical similarity of word referents was greater in BA 19 than in BA 17 (t(26) = 2.62, p = 0.045, Cohen’s d = 0.5), with no significant effects in other comparisons (both p values > 0.1). In the blind participants, no differences across the visual areas were found (all p values > 0.25). In line with the results of the main analysis, none of the above-described tests produced significant effects when the correlations with the ratings of conceptual similarity of word referents were considered (all p values > 0.15). This further suggests that activity patterns in the visual areas do not reflect the conceptual similarity between word referents in either sighted or blind individuals.

As a control analysis, we searched for the representation of more abstract, conceptual similarity between word referents in the visual areas in the left hemisphere only, instead of using bilateral masks. While the responses to linguistic stimuli in the blind brain are less lateralized than in the sighted brain (Lane et al., 2017; Dzięgiel-Fivet and Jednoróg, 2024), one can perhaps still argue that the representational plasticity for language in blind individuals should be more prominent in the left hemisphere. Contrary to this argument, our analysis in the left hemisphere produced results that are virtually identical to those described above (Fig. S2).

Furthermore, we asked whether conceptual similarity between word referents is computed in the left anterior temporal lobe (ATL), which is involved in the processing of abstract knowledge in both sighted and blind individuals (Striem-Amit et al., 2018; Wang et al., 2020). Indeed, we observed significant correlations between similarity in activity patterns for words in this region and the ratings of conceptual similarity of word referents in both groups (both p values < 0.05) (Fig. S3). The 2 (group) x 2 (similarity metric) x 2 (brain area) ANOVA, which compared the correlation scores for the visual cortex and the ATL, indicated a significant interaction between the similarity metric and the brain area (F(1,45) = 7.85, p = 0.007, partial eta squared = 0.15). No other main effects or interactions were significant. The post-hoc tests showed that the correlation with conceptual similarity ratings were higher in the ATL than in the visual cortex (p = 0.038), whereas the correlation with physical similarity ratings were higher in the visual cortex than in the ATL (p = 0.008). Furthermore, the correlation with the physical similarity ratings was significantly higher than the correlation with the conceptual similarity ratings in the visual cortex (p = 0.006), but not in the ATL (p > 0.25).

Finally, we directly compared the neural similarity matrices reflecting the similarity in activity patterns for specific words in the visual areas across the blind and the sighted participants (Fig. 4). To this aim, we correlated such a neural similarity matrix of each participant with the analogous neural similarity matrix of every participant from the other group (sighted-blind, blind-sighted). The obtained correlation scores were then compared to the within-group correlations (sighted-sighted, blind-blind) produced in the analogous way.

**Figure 4.**
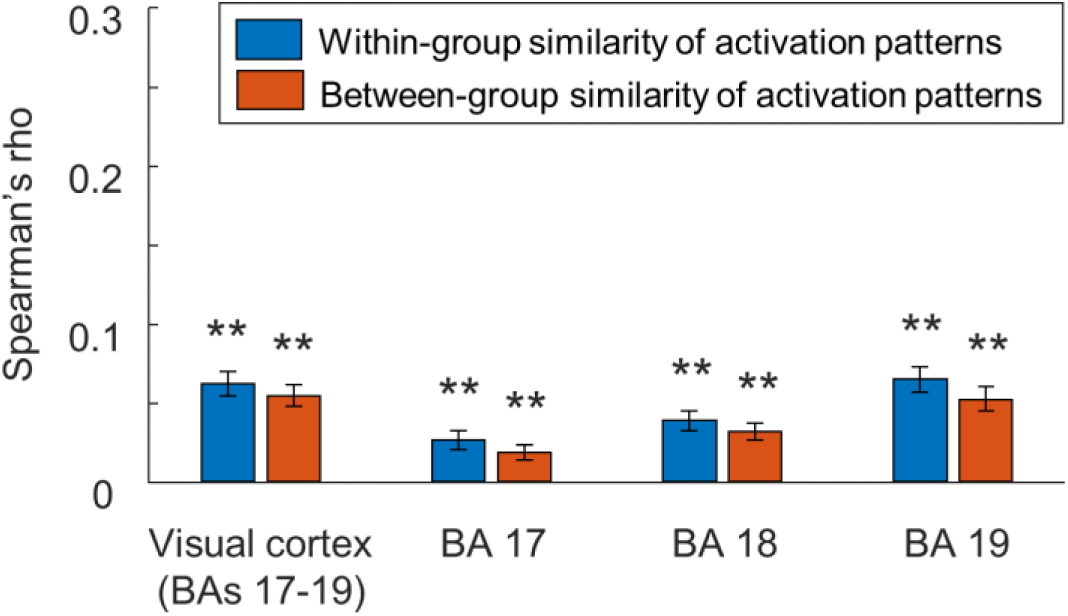
Between-group similarity in activity patterns for words in the visual areas of blind and sighted participants is comparable to within-group similarities. The between-group similarity scores were calculated by correlating the neural similarity matrices of each participant, reflecting similarity between the activity patterns for specific words in the visual areas, with the neural similarity matrices of every participant in the other group (blind-sighted, sighted-blind). The within-group similarity scores were calculated by correlating the neural similarity matrices of each participant with the neural similarity matrices of every other participant from a given group (blind-blind, sighted-sighted). ** p < 0.01, corrected for multiple comparisons using the false discovery rate. Error bars represent the standard error of the mean.

If word processing induces different types of neural representations in the visual areas of sighted and blind individuals, then the between-group correlations should be either non-significant or markedly lower than the within-group correlations. However, we observed robust between-group correlations in the visual cortex (BAs 17, 18, and 19 combined; p < 0.01) and in all specific visual areas (all p values < 0.01). Moreover, we did not find significant differences in the between-group and the within-group correlation scores in any of the visual regions (all p values > 0.1). The 2 (type of correlation) x 3 (visual area) ANOVA produced only a significant main effect of the visual area (F(2,92) = 25.41, p < 0.001, partial eta squared = 0.37), indicating that both types of correlations linearly increased from BA 17 to BA 19 (all p values < 0.001, linear contrast: p < 0.001). Neither the main effect of the type of correlation nor the interaction were significant (both p values > 0.15). These results further support our hypothesis and suggest that representations activated by words in the visual areas of blind and sighted individuals are related.

### Representational connectivity of the visual cortex

Next, we investigated the “representational connectivity” (Kriegeskorte et al., 2008) of the visual cortex in blind and sighted participants during word processing (Fig. 5). That is, we searched for brain regions in which words induce similar activity patterns to those induced in the visual cortex. The aim of this analysis was to investigate whether the position of the visual cortex in the cortical processing hierarchy during word processing is comparable in blind and sighted individuals. To study this issue, we correlated the neural similarity matrices reflecting the similarity in activity patterns for specific words in the visual cortex with the analogous neural similarity matrices obtained for every other Brodmann area found in the BrainVoyager cortical atlas (33 target areas in total; see Methods). To account for effects that are present across the whole cortex, we calculated the average correlation between the visual cortex and the target areas and subtracted this score from the results. This allowed us to identify brain regions in which the word-induced representations are more or less similar than the cortical average to the representations computed in the visual cortex.

**Figure 5.**
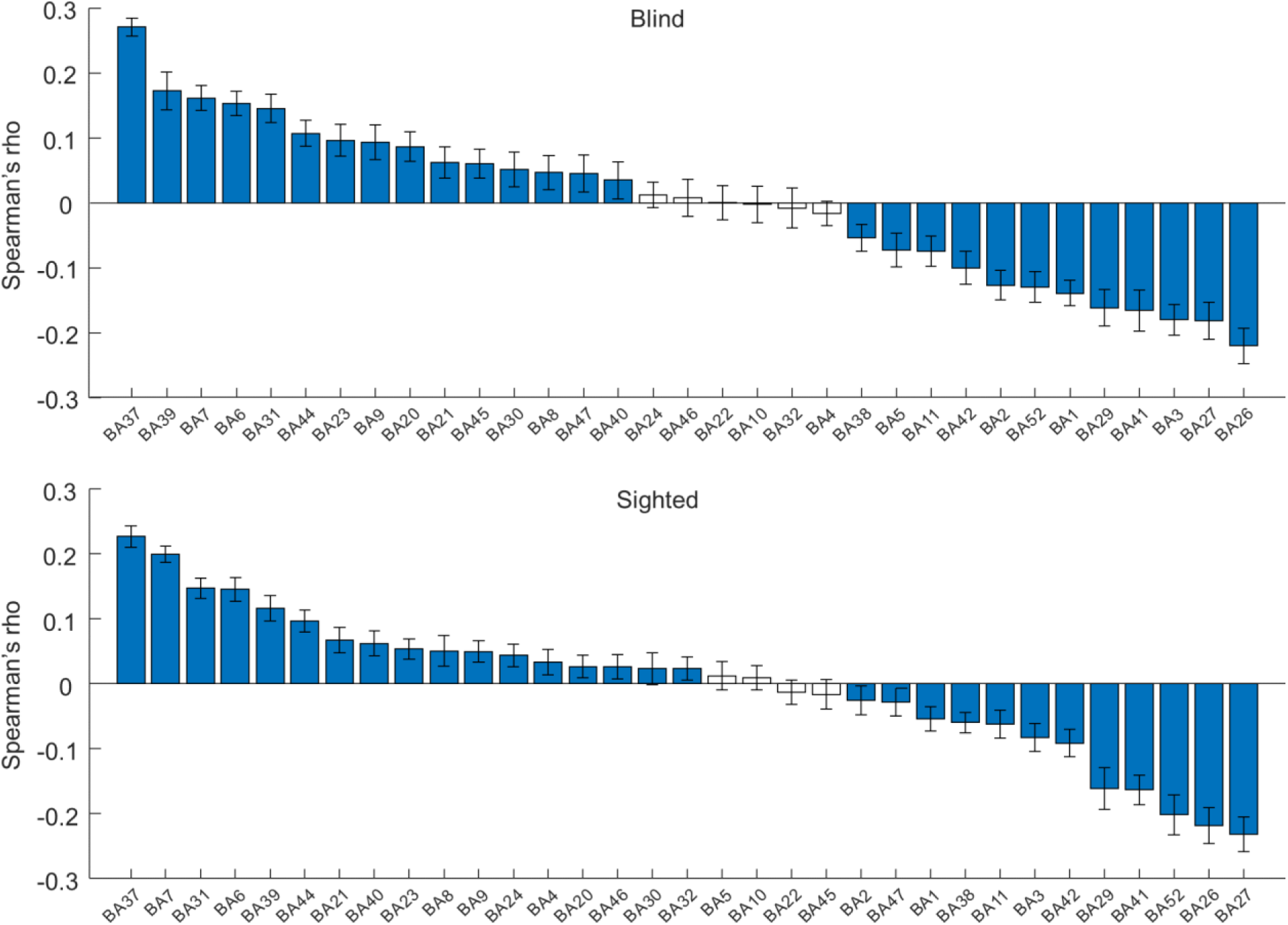
During word processing, the visual cortex in both blind and sighted participants shows the greatest “representational connectivity” to the occipitotemporal cortex. For each brain region, the representational connectivity to the visual cortex was calculated by correlating neural similarity matrices, reflecting similarity between the activity patterns for specific words, obtained for the visual cortex and this region. In each participant group, the average correlation between the visual cortex and the target brain regions was subtracted from the results, to visualize regions that show higher-than-average and lower-than-average representational connectivity to the visual cortex during the word processing. The results for such regions are marked in blue (comparison against average correlation, p < 0.05, corrected for multiple comparisons using the false discovery rate). The target brain regions were sorted by the obtained representational connectivity scores. Error bars represent the standard error of the mean.

In both participant groups, the visual cortex showed the greatest representational connectivity to the occipitotemporal cortex (BA 37). The representational connectivity between these two adjacent regions was higher than the representational connectivity between the visual cortex and its every other adjacent region in the blind participants (BAs 7, 30, 31, and 39; all p values < 0.002), and every other adjacent region, except for BA 7, in the sighted participant (all p values < 0.001). Thus, the preferential representational link between the visual cortex and the occipitotemporal cortex cannot be explained merely by the topographical proximity of these two regions. Conversely, the visual areas in both groups showed lower-than-average representational connectivity to primary auditory and somatosensory cortices (BAs 41-42 and BAs 1-3, respectively). Finally, the representational connectivity scores obtained in both participant groups were highly correlated (Spearman’s rho = 0.91, p < 0.001), which further suggests a similar position of the visual cortex in the cortical processing hierarchy for word processing in these two groups. Virtually the same results were observed when only the lower-level visual areas (BA 17 or BA 18) were included in the analysis (Figs. S4-5).

### Searchlight analysis

We performed a searchlight analysis to investigate what brain regions, beyond the visual cortex, are involved in representing physical and conceptual similarity between word referents (Fig. 6). First, we ran the analysis with participants from both groups combined in one sample, to achieve the greatest statistical power to reveal the common effects (the results for each group separately are presented in Fig. S6). Second, we compared the results across groups to visualize differential effects. As in the analysis in the visual areas, the partial correlation approach was used to account for the common variance in the ratings of physical and conceptual similarity.

**Figure 6.**
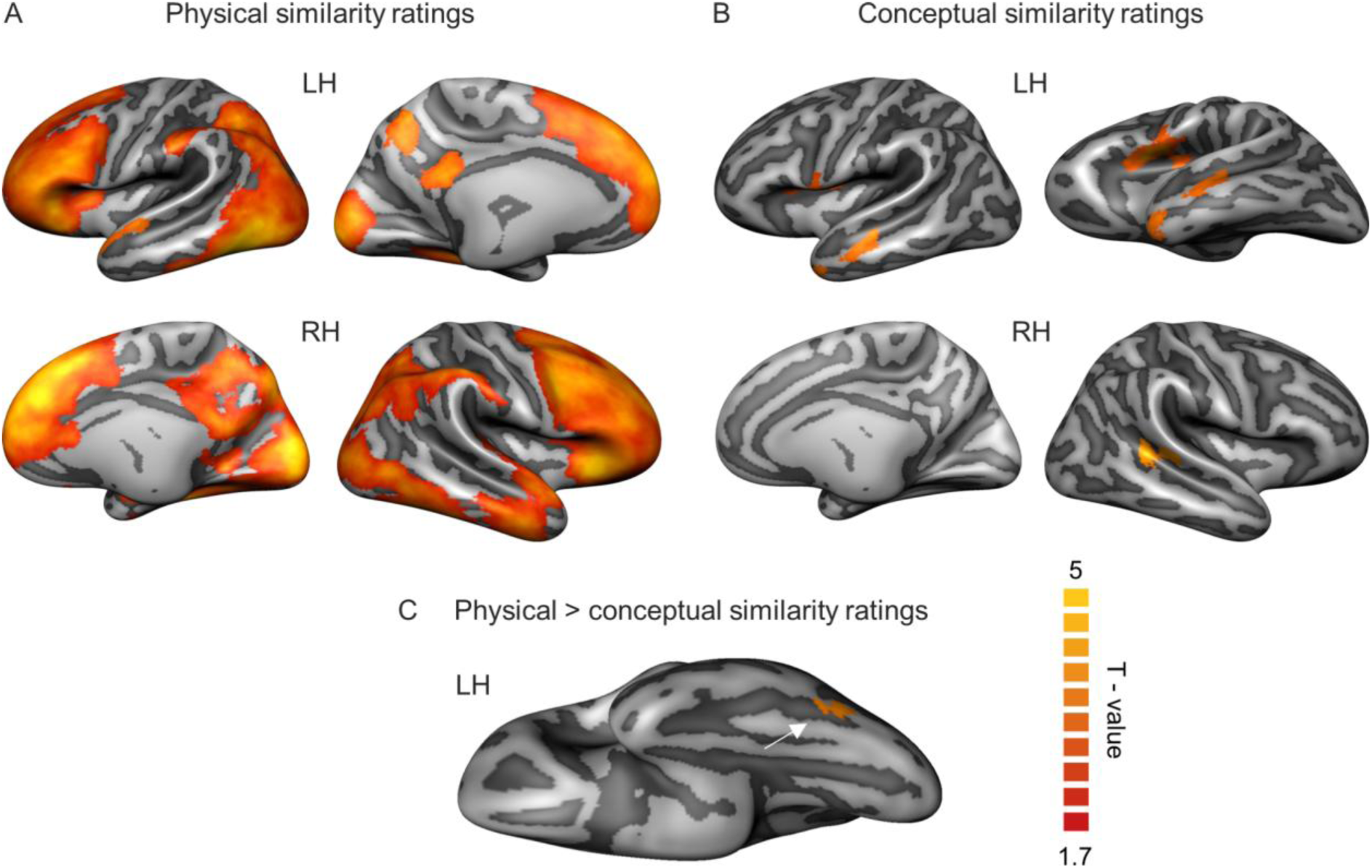
Searchlight analysis. (A-B) The results of correlation between the neural similarity matrices, reflecting similarity between the activity patterns for specific words, and the behavioral ratings of (A) physical and (B) conceptual similarity between the word referents. (C) Brain regions in which the correlations with physical similarity ratings were significantly higher than the correlations with conceptual similarity ratings. The white arrow points to the effect in the occipitotemporal cortex. The analyses combined blind and sighted participants in one sample, to achieve the greatest statistical power to detect common effects. Between-group comparisons did not produce significant results. In all analyses, The statistical significance was determined using threshold-free cluster enhancement (TFCE) maps and Monte Carlo simulation. The statistical threshold was set at p < 0.05, corrected for multiple comparisons. The T-values for the effects that survived the threshold are presented for the visualization purposes.

When all participants were combined in one sample, the ratings of physical similarity of word referents correlated with similarity in activity patterns for words in the canonical language and semantic regions, such as the frontal cortex, the anterior temporal lobe, and the precuneus (Fig 6A). Furthermore, significant effects were found throughout the visual cortex and in the lateral and ventral occipitotemporal regions. The analogous analysis including the ratings of conceptual similarity of word referents produced more constrained, but robust effects in the left anterior temporal lobe, the superior temporal sulci, the left middle temporal gyrus, the left frontal operculum, and the left insula (Fig. 6B). No significant effects were detected in the visual areas. The direct comparison between the results of these two analyses showed significantly stronger correlation with the physical similarity ratings, compared to the conceptual similarity ratings, in the left ventral occipitotemporal cortex, at the border of BAs 19 and 37 (Fig. 6C). No differences between groups and no interaction between the group and the similarity metric were observed, even at the level of statistical trend (tested at p < 0.1, corrected for multiple comparisons).

### Univariate analysis

We also performed the univariate analysis, to investigate the magnitude of activations for spoken and written words in the visual areas in both groups (Fig. 7). We particularly asked whether the processing of spoken and Braille words activated the visual cortex in the blind participants, as was reported previously.

**Figure 7.**
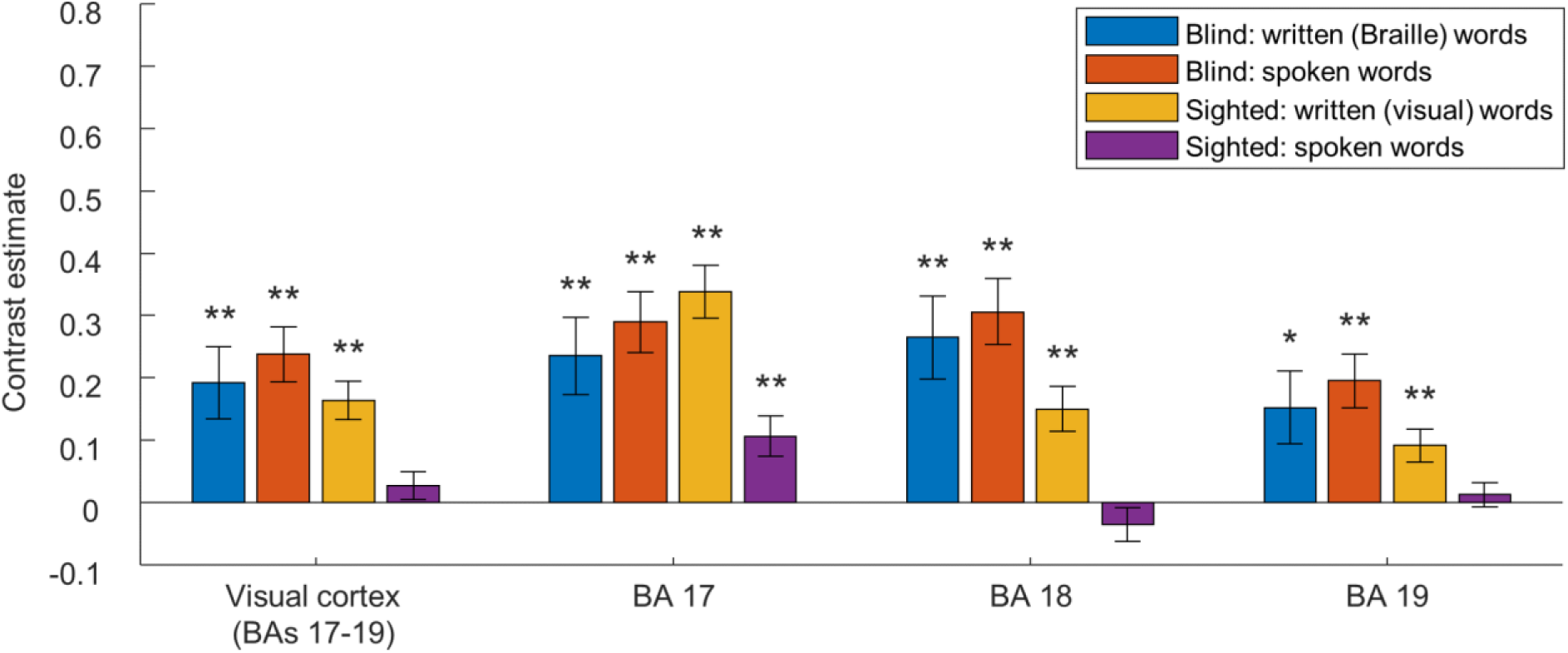
Univariate analysis of activations in the visual cortex. The average activations for spoken and written words, compared to rest periods, in the visual areas of blind and sighted participants. * p < 0.05, ** p < 0.01, corrected for multiple comparisons using the false discovery rate. Error bars represent the standard error of the mean.

Indeed, we found that, compared to rest periods, both spoken and written (Braille) words robustly activated the visual cortex and all specific visual areas in the blind participants (all p values < 0.05) (Fig. 7A). In the sighted participants, all visual areas were activated by written (visual) words (all p values < 0.01). The spoken words activated the primary visual cortex (BA 17; p = 0.002), but not the higher-level visual areas (p > 0.25), in this group (see also: Seydell-Greenwald et al., 2023). The 2 (group) x 2 (word presentation modality) x 3 (visual area) ANOVA produced a significant three-way interaction (F(2,90) = 15.73, p < 0.001, partial eta squared = 0.26). The comparison across the groups confirmed that, in all visual areas tested, activations for spoken and written (Braille) words in the blind participants were stronger than the activations for spoken words in the sighted participants (all p values < 0.05). The comparisons across the modalities showed that, in the blind participants, spoken and written word presentations induced comparable activations in all visual areas (all p values > 0.25). As could be expected, in the sighted participants, activations for written (visual) words were stronger than the activations for spoken words (all p values < 0.05). Finally, the comparison across the visual areas revealed that, in the blind group, activations for both spoken and written words were stronger in lower-level visual areas (BAs 17 and 18) than in the higher-level visual areas (BA 19) (all p values < 0.059). In the sighted participants, the activation in the primary visual cortex (BA 17) was stronger than the activation in the higher-level visual areas (BAs 18 and 19), again for both modalities of word presentation (all p values < 0.001).

The whole-brain analysis (Fig. S7-S8) confirmed the pattern of results described above. Additionally, this analysis revealed that, in both groups, both modalities of word presentation induced expected activations in the canonical language network. Apart from decreased activation for Braille words in the cuneus and the precuneus in the blind groups, we did not find any significant differences between groups or the modalities of word presentation in the canonical language or semantic regions.

### Influence of word presentation modality on the results

Previous studies with blind participants reported the representation of physical properties of word referents in the occipitotemporal cortex (Mahon et al., 2009; He et al., 2013; Peleen et al., 2013, 2014; Xu et al., 2023). Why did these studies not find the effects in the occipital lobe, which we robustly detected in our study? One explanation is that these studies used only the spoken word presentation. The activity of the visual areas is modulated by auditory stimulation (Laurienti et al., 2002; Iurilli et al., 2012; Murray et al., 2016; Anurova et al., 2019; Ferraro et al., 2020). Sounds with different acoustic properties induce different activity patterns in the low-level visual areas (Martinelli et al., 2020). These modulations of activity in the visual cortex, by the auditory cortex, could make effects related to the processing of word meaning less discernible. In line with this hypothesis, studies that found the representation of physical properties of word referents in the occipital lobe of sighted individuals used visual rather than auditory word presentation (Borghesani et al., 2016; Martin et al., 2018).

To explore this possibility in our data, we divided them into runs with spoken word presentation and written word presentation. We then compared the correlations between the similarity in activity patterns for words and the ratings of physical and conceptual similarity of word referents in these two datasets (Fig. S9). Indeed, the results for written words were stronger in both groups. The 2 (group) x 2 (similarity metric) x 3 (visual area) x 2 (modality of word presentation) ANOVA produced a significant main effect of modality of word presentation (F(1,45) = 5.21, p = 0.027, partial eta squared = 0.1), indicating that the correlation scores in the visual cortex were generally higher for written words. Furthermore, we also found the main effect of similarity metric (F(1,45) = 4.3, p = 0.044, partial eta squared = 0.09), indicating that our key finding - significant correlation of the activity patterns for words in the visual cortex with physical, but not conceptual similarity between the word referents in both groups - replicated also in this additional analysis.

## Discussion

In this study, we found that visual areas of both congenitally blind and sighted participants represented differences between individual concrete words in a format independent of the word presentation modality. We further observed that, in both participant groups, the activation patterns for words in the visual cortex reflected physical, but not more abstract, conceptual similarity between word referents. Subsequently, we directly compared activation patterns induced by words in the visual areas of blind and sighted participants. We found that the between-group correlations in these activity patterns were comparable to the within-group correlations. Finally, we demonstrated that, during word processing, the visual cortex in both blind and sighted participants showed the greatest “representational connectivity” to the same cortical region, the occipitotemporal cortex. All these between-group similarities were observed despite the fact that, as in previous studies, the processing of spoken and Braille words by the blind participants induced markedly stronger responses in the visual cortex than the processing of spoken words by the sighted participants.

Our study shows that, during word processing, the activation patterns in visual areas of congenitally blind and sighted individuals are related and reflect a specific semantic dimension - the physical similarity between word referents. Previous studies have repeatedly demonstrated that processing of linguistic stimuli, such as words and sentences, strongly activates the visual areas in blind individuals, in the absence of comparable results in sighted participants (Sadato et al., 1996; Burton et al., 2002; Bedny et al., 2011; Bedny et al., 2015; Lane et al., 2015). An increasingly popular interpretation of these findings is that, in the absence of visual inputs, the blind visual cortex assumes higher cognitive functions, such as processing of abstract semantic or grammatical information, which are not processed by the visual areas of the sighted (Bedny, 2017; Saccone et al., 2024). Our results suggest a more parsimonious explanation: responses to linguistic stimuli in the blind visual cortex can be driven by representational mechanisms present also in the visual areas of sighted adults. These mechanisms might produce stronger activations in the blind visual cortex because, in the absence of visual inputs, this region develops weaker inhibitory pathways (Benevento et al., 1995; Morales et al., 2002) and supranormal sensitivity to stimulation (Merabet et al., 2008; Binda et al., 2018). In other words, our results suggest that linguistic stimuli, such as words, activate comparable neural representations in the visual areas of blind and sighted individuals, but this process results in stronger activation of the blind visual cortex due to its enhanced excitability.

Consequently, our findings suggest that responses to language in the blind visual cortex can be driven by representational mechanisms that support visual perception in the sighted brain. In the visual areas of sighted individuals, the processing of visual feedforward signals is modulated by predictions generated in the higher-level brain regions (Rao and Ballard, 1999; Lee and Mumford. 2003; Friston, 2005). Our results suggest that such modulatory and potentially predictive signals can be generated even based on symbolic stimuli, such as words (see also: Seydell-Greenwald et al., 2023). Based on our results, one might hypothesize that language comprehension activates a top-down informational cascade, in which the knowledge about word referents is first backprojected from the canonical language regions to richly multimodal shape- and size-processing areas in the occipitotemporal cortex (Ricciardi et al., 2014; Bi et al., 2016). These areas, in turn, might communicate the physical properties of word referents, most useful for visual predictions, to the occipital lobe. Our findings suggest that such a neural architecture of backprojections can develop and function even in the absence of visual experience.

Previous studies documented the representation of the physical properties of word referents in the visual areas of sighted individuals (Borghesani et al., 2016; Martin et al., 2018). In blind individuals, such representations were found primarily in the occipitotemporal cortex (Mahon et al., 2009; He et al., 2013; Peleen et al., 2013, 2014; Xu et al., 2023), with one recent study showing that, in the lateral occipitotemporal cortex, the representation of physical knowledge is not accompanied by representations of more abstract properties in either blind or sighted individuals (Xu et al., 2023). Here, we document the representation of physical, but not abstract properties of word referents throughout the visual cortex of blind individuals. Interestingly, in sighted participants, correlations with the ratings of physical similarity of word referents were significant in the associative and secondary visual areas, but not in the primary visual cortex (BA 17). In line with this result, the correlation between the activations patterns in the primary visual cortex and the ratings of physical similarity of word referents was significantly lower in the sighted group than in the blind group. This between-group difference was observed even though the other analyses showed that the representation of differences between words in the primary visual cortex is of comparable precision in both groups (Fig. 2), and that activation patterns induced by words in this area are related across groups (Fig. 4). One explanation of these results is that the behavioral ratings of physical similarity might predominantly capture the global physical properties of word referents, such as overall shape or size. One can speculate that, in sighted individuals, the primary visual cortex is better suited to represent lower-level features, such as spatial frequencies, contrasts, or orientations (Morgan et al., 2019). In blind macaques, the projections from high-level ventral visual areas to the primary visual cortex are strengthened, and the average length of intrinsic connectivity in parts of the primary visual cortex is increased (Magrou et al., 2018). Potentially, similar anatomical plasticity might support representations of more global physical properties in the primary visual cortex of blind humans. Such representation might still be related to lower-level representations in the sighted early visual areas, but, at the same time, might be better described by the behavioral ratings of physical similarity. Overall, our study shows that word processing activates related representations of physical knowledge in the visual cortex of both congenitally blind and sighted individuals. This, however, does not preclude a certain degree of plastic changes in these representations in the blind visual cortex.

The semantic mechanism reported in our study can explain a number of effects found in the blind visual cortex for linguistic stimuli. The gradual increase in activation magnitude for pseudowords, words, and sentences (Bedny et al., 2011) can be explained by progressively richer information about the physical environment conveyed by these stimuli (compare “an allb” vs. “a ball” vs. “Mary threw a ball to the dog”). Decoding of word semantic categories (Abboud et al., 2019) might rely on systematically different physical properties of word referents across these categories. Stronger activation of the visual cortex during a semantic task, compared to a phonological task (Burton et al., 2003), might indicate increased salience of the physical representations of word referents when one thinks about word meaning. Arguably, however, the proposed mechanism is unlikely to be the only source of language-related activations in the visual areas of blind individuals. Importantly, our study directly compared the activation patterns induced by words in the visual areas of sighted and blind individuals (Fig. 4). This analysis was unconstrained by any theoretical model, therefore, it should capture the influence of every major input to the visual cortex on the representations that emerge in this region during word processing. The results showed that these representations, computed in the visual cortex of sighted and blind individuals, are related. This further suggests that, in both populations, the emergence of these representations is driven by similar mechanisms.

Our findings contribute to a better understanding of principles of functional plasticity in the human brain. One way to think about activations for language in the blind visual cortex is that, in the absence of visual signals, this region takes on new cognitive functions, which are radically different from those computed in the sighted visual cortex (Bedny et al., 2017; Saccone et al., 2024). Our study provides a different perspective and suggests that responses to language in the blind visual cortex can be driven by representational mechanisms that are present in the sighted adult brain and are uncovered in the absence of visual inputs (Pascual-Leone and Hamilton, 2001; Makin and Krakauer, 2023). We have already made that argument in our recent work, in which we demonstrated that the motion-sensitive area V5/MT in blind individuals shows different activity patterns for concrete nouns and verbs, in the absence of significant results for abstract and pseudo nouns and verbs (Urbaniak et al., 2024). Based on these results, we suggested that this region represents motion connotations of noun and verb referents, more salient in the concrete word category, rather than more abstract grammatical or conceptual distinctions, present in all word categories. This suggests that, in blind individuals, this visual region might represent linguistic stimuli through physical features of word referents, retrieved from semantic representations. Here, we show that this principle may apply throughout the visual cortex of blind individuals, even in the early visual areas, which have been proposed to be a hotspot of functional plasticity in the blind brain (Amedi et al., 2003). Moreover, we directly demonstrated that processing of words induces the same type of representations in the blind and the sighted visual cortex, a result that validates a critical prediction of our hypothesis.

Furthermore, our results provide insight into mechanisms that underpin top-down modulations of the activity in the visual cortex. The nature of such top-down modulatory signals - particularly their relationship to visual imagery - is still debated. Our study documents the emergence of similar representations of word referents in visual areas of sighted individuals and in visual areas of congenitally blind individuals, who could not develop visual imagery. Previous studies have already shown compelling parallels in the functional organization of the occipital lobe in sighted and blind individuals (Pascual-Leone and Hamilton, 2001; Ricciardi et al., 2014; Heimler et al., 2015). These studies, however, mostly presented blind participants with sounds or shapes, which makes the effects related to lateral projections from the other sensory systems and the backprojections from the higher-level brain regions difficult to disentangle. Our work, in contrast, documents the emergence of semantic representation in the visual areas of both populations, a process that cannot be explained by lateral interactions between sensory systems. This finding strongly suggests that the functional modulation of the visual cortex by backprojections from the higher-level regions is a process that is independent of visual imagery (see also: Seydell-Greenwald et al., 2023). An intriguing hypothesis is that the functional modulations of the visual cortex, akin to those observed in our study, constitute a mechanism that is evolutionarily older than the conscious, internally generated visual imagery. One can suppose that backprojections to the visual cortex, which originally emerged in the brain to support visual perception, were then utilized for more general visuospatial processing, such as visual imagery or visual working memory, ultimately becoming the brain’s “spatial blackboard” (Roelfsema and de Lange, 2016; Linton, 2021). It remains to be tested whether this architecture can causally support spatial thinking or spatial memory in blind individuals.

The idea that functional plasticity in the visual cortex of blind individuals “unmasks” processes and computations present also in the sighted visual cortex has already received considerable support from studies of auditory and tactile processing (Pascual-Leone and Hamilton, 2001; Heimler et al., 2015). However, the discovery of activations for words and sentences in the blind visual cortex has been sometimes treated as prime evidence against this hypothesis (Bedny et al., 2017). Our study counters this argument and suggests that enhanced activations of the blind visual cortex during auditory, tactile, and semantic processing may be driven by similar mechanisms.

In summary, we found that, during word processing, the activation patterns in visual areas of congenitally blind and sighted individuals are related and reflect a specific semantic dimension - the physical similarity between word referents. This shows that responses to linguistic stimuli in the blind visual cortex can be driven by representational mechanisms that are present in the sighted adult brain and are uncovered in the absence of visual inputs. Our study suggests that the type of neural representations computed in the visual cortex cannot be easily changed, even by dramatic changes in visual experience.

## Methods

### Participants

Twenty congenitally blind subjects (9 males, 11 females, mean age = 36.4 y, SD = 7.61 y, average length of education = 15.2 y, range = 12 - 17 y) and 27 sighted subjects (12 males, 15 females, mean age = 35.7 y, SD = 8.6 y, average length of education = 15.56 y, range = 12 - 17 y) participated in the study. Two participants in the blind group and three in the sighted group were left-handed, while the remaining participants were right-handed. The blind and the sighted groups were matched for age, gender, handedness, and length of education (Mann-Whitney and Chi-square tests, all p values > 0.25). In the blind group, blindness had a variety of causes, including the retinopathy of prematurity, retinal detachment, atrophy of the optic nerve, or unknown causes. Nine blind participants reported to have some light perception, but no object or contour vision. Additionally, two blind participants reported to have some form of contour vision, which, however, was not precise enough to be functional. All blind participants were proficient Braille readers. All blind and sighted participants were native Polish speakers, had normal hearing, and had no history of neurological disorders, apart from the cause of blindness in the blind group. All participants had no contraindications to the MRI, gave written informed consent and were paid for participation. The study was approved by the ethics committee of Institute of Psychology, Polish Academy of Sciences.

### Stimuli

Twenty Polish nouns referring to concrete entities (e.g., “a cup” or “a snake”; see Fig. S1 for a complete list) were used as stimuli. The words had from 3 to 5 letters. The length restriction was introduced because the Braille display used in the fMRI study (see below) can display up to 5 letters at the same time. The words’ frequency of occurrence in Polish language was moderate to high (range of Zipf frequency values = 2.99 - 4.76, as indicated by the Subtlex-pl database; Mandera et al., 2015). The word list was inspired by the words used by Martin et al. (2018). Their list was shortened and adjusted to meet requirements listed above.

### fMRI experiment

In the fMRI experiment, the participants were asked to either listen to or read the words and answer the questions about either physical or conceptual properties of word referents. In half of the fMRI experiment, the words were presented in the spoken modality, whereas, in the other half, in the written modality (in the visual alphabet in the sighted group and in the Braille alphabet in the blind group). The modality of word presentation was alternated across fMRI runs in an odd / even scheme. There were 12 runs (6 spoken and 6 written runs).

In each of the runs, there were two experimental blocks - in one of them the participants were asked to think about physical properties of word referents, whereas, in the other, about their more abstract, conceptual properties. Each block started with a 2.5-s presentation of a question about a specific physical (e.g., is this elongated?) or conceptual (e.g., is this animate?) property of word referents. Then, after 12-s break, all 20 words were presented, one after another. The participants’ task was to answer the question for each word referent by providing a “yes” or “no” answer. The blocks were separated by 12-s breaks, and there were 4 s of rest at the beginning and at least 16 s of rest at the end of each run.

In the sighted participants, the questions were presented in either the spoken or the written modality - the modality of question and word presentation were matched in each run. In the blind participants, such a solution was not practical because of the maximal capacity of the Braille display we used during fMRI and a relatively slower pace of Braille reading, compared to visual reading. Thus, in the blind group, the questions were always presented in the spoken modality. Furthermore, the duration of word presentation was also adjusted to the needs of each presentation modality - each word was presented for 1 s in the spoken modality (both groups) and in the visual alphabet (the sighted group only), and for 2.5 s in the Braille modality (blind group only). In each presentation modality, every word presentation was followed by a rest period lasting for either 2 s (67% probability) or 6 s (33% probability). Such a pace was sufficient to ensure word comprehension in all sighted and blind pilot participants. The post-fMRI interviews confirmed that blind and sighted individuals participating in the actual experiment found this pace comfortable.

In the sighted group, the spoken and the written runs had the same length (approx. 4 mins and 4 s) and the modality of a first run was randomly assigned to each participant. In the blind group, the spoken runs were of the same length, but written runs were longer (approx. 4 mins and 43 s) because of longer word presentations in the Braille modality. Because of different run lengths, the modality of the first run had to be fixed for all participants, in order to avoid errors during the data collection - thus, in the blind group, all participants started from the spoken run.

There were 6 different questions about physical properties of word referents and 6 different questions about their conceptual properties (Table 1). Each question was presented twice during the experiment - once in the spoken run and once in the written run. The order of blocks with physical questions and conceptual questions was counterbalanced within each participant and each word presentation modality - that is, in the first half of the experiment, the runs started from one attentional condition, whereas, in the second half, from the other attentional condition. What condition was first in the first and the second half of the experiment was randomly drawn for each participant.

**Table 1.** English translations of the questions used in the fMRI experiment.

| Questions about physical properties | Questions about conceptual properties |
| --- | --- |
| Is this round? | Is this natural? |
| Is this angular? | Is this manufactured? |
| Is this smooth? | Is this living? |
| Is this rough? | Is this non-living? |
| Is this elongated? | Is this pleasant |
| Is this symmetrical? | Is this unpleasant? |

The stimuli presentation was controlled by a program written in PsychoPy v2022.2.3 (Peirce, 2007). The sounds were presented through MRI-compatible headphones. The visual alphabet was presented on a screen (white letters on a gray background), which participants watched in a mirror attached to the MRI head coil. The Braille alphabet was presented through a custom-made, MRI-compatible Braille display (Debowska, 2013), which was strapped to the left or right thigh of a participant so that it could be comfortably reached by the hand that this participant usually uses for Braille reading. The responses were provided by a two-button response pad. Before starting the experiment, each participant completed a short training session, to ensure that the written words are visible / reachable. Furthermore, the volume of spoken word presentation was individually adjusted to a loud, but comfortable level.

### Behavioral rating experiment

The ratings of physical and conceptual similarity between the word referents were obtained through an online experiment. Four hundred and two native Polish speakers (200 males, 202 females, mean age = 28.9 y, SD = 9 y) participated in this study. Each participant was visually presented with 30 word pairs that were randomly drawn from the set of 190 word pairs that could be created from the 20 words used as stimuli. For each pair, the participants were instructed to rate physical similarity of word referents, while ignoring conceptual similarity, and conceptual similarity, while ignoring physical features. In rating the physical similarity, the participants were instructed to focus on properties such as shape, size, and color. They were given an example of “a ball” and “an apple” as words naming physically similar objects. In rating the conceptual similarity, the participants were instructed to focus on their knowledge about word referents, their function, where they can be found, and similar properties. They were given an example of “an apple” and “a banana” as words naming conceptually similar objects. The participants saw a word pair, rated the physical and conceptual similarity of word referents on 5-point scales, and confirmed their answers with a keyboard press in order to see a next word pair.

The experiment was prepared in Psychopy v2022.2.3 and uploaded to Pavlovia online repository (https://pavlovia.org). The participants were recruited through the Prolific platform (https://www.prolific.com) and were paid for the participation. The results obtained for each word pair were averaged across the participants. The obtained matrices of average ratings were then used in the analysis of neuroimaging data.

### Neuroimaging parameters

The neuroimaging data were acquired on a 3T Siemens Magnetom Prisma Fit MRI scanner using a 32-channel head coil at the Laboratory of Brain Imaging in the Nencki Institute of Experimental Biology in Warsaw. Functional data were acquired using a multiband sequence with the following parameters: 60 slices, phase encoding direction from posterior to anterior; voxel size: 2,5 mm^3^; TR = 1.41 s; TE: 30.4 ms; multiband factor: 3. Before the start of the first functional run, T1-weighted anatomic scans were acquired using MPRAGE sequence with the following parameters: 208 slices, phase encoding direction from anterior to posterior; voxel size: 0,8 mm^3^; TR = 2.5 s; TE: 21.7 ms.

### Neuroimaging data analysis

#### Preprocessing

The neuroimaging data were converted from the DICOM to the NIFTI format using the dcm2niix (Li et al., 2016). Then, the data were preprocessed in BrainVoyager 22.4 (BrainInnovation). Standard routines were used to preprocess the functional data, including slice scan time correction, 3D rigid body motion correction, and temporal high-pass filter (GLM with Fourier basis set, 2 cycles per run). For each participant, the functional data were mapped onto an individual reconstruction of the cortical surface, created based on the collected anatomical image. All subsequent preprocessing steps and analyses were performed in the surface space. The functional data were spatially smoothed for the univariate analysis (the nearest neighbors approach, repeat value: 4). No spatial smoothing was performed for the multi-voxel pattern analysis.

Two first-level statistical models were created for each subject. For the multi-voxel pattern analysis, the data were modeled at the level of single words (20 words x 2 experimental blocks in each run, 40 predictors per run). For the univariate analysis, the data were modeled at the level of word blocks, understood as time between the onset of the first word in a given experimental block and the offset of the last word in this block (2 predictors per run). In both models, questions that were presented before the word blocks were additionally modeled as conditions of no interest (2 predictors per run). Signal time course was modeled using a general linear model (Friston et al., 1995) by convolving a canonical hemodynamic response function with the time series of predictors.

#### Multi-voxel pattern analysis

All multi-voxel pattern analyses were performed in CosmoMVPA (v.1.1.0; Oosterhof et al., 2016), running on Matlab R2022b (MathWorks). The NeuroElf toolbox (https://neuroelf.net/) was used to import BrainVoyager files into CosmoMVPA. All analyses were performed on T-values reflecting activation for each word presentation relative to average activation during rest periods in a given run (Misaki et al., 2010). Thus, 480 T-maps (20 words x 2 experimental blocks x 12 runs) were calculated and included in the analysis for each participant.

The multi-voxel pattern classification analyses were performed using a linear support vector machine classification algorithm, as implemented in the LIBSVM toolbox (v. 3.23; Chang and Lin, 2001). A standard LIBSVM data normalization procedure (i.e., Z-scoring beta estimates for each voxel in the training set and applying output values to the test set) was applied to the data before classification. The classification was performed in an odd-even cross validation scheme, that is, the classifier was trained on the odd runs and tested on the even runs, and vice versa. As a result, the algorithm was always trained on one sensory modality of the word presentation and tested on the other.

The representational similarity analyses were performed on brain activations (quantified as T-values, see above) that were averaged across runs and experimental blocks for each word. For each participant, the neural similarity matrices were calculated by correlating activity patterns for words in a given area with each other using Pearson correlation. Then, these matrices were correlated with the behavioral ratings of physical and conceptual similarity between word referents using Spearman’s correlation. Partial correlation approach, as implemented in CosmoMVPA, was used to account for common variance in the physical and conceptual similarity ratings.

The neural similarity matrices obtained in the visual cortex and in specific visual areas were correlated across the participants, to obtain the between-group and the within group similarity scores. To calculate the between-group similarity score, the neural similarity matrices of each participant were correlated with the neural similarity matrices of every participant from the other group (sighted-blind, blind-sighted). These values were then averaged to obtain one score for each participant. These scores were compared with within-group similarity scores, which were obtained by correlating the neural similarity matrices of each participant with the neural similarity matrices of every other participant from the same group (sighted-sighted, blind-blind). Spearman’s correlation was used to obtain all these correlation scores.

Finally, to obtain the “representational connectivity” scores (Kriegeskorte et al., 2008), the neural similarity matrices obtained in the visual areas of each participant was correlated with neural similarity matrices obtained for this participant in all other Brodmann areas found in the BrainVoyager cortical atlas (see below; 33 target areas in total). To account for effects present across the whole cortex, we calculated the average correlation across all target areas and subtracted this score from the results. This allowed us to identify brain regions in which the word-induced representation is more or less similar than the cortical average to the representation computed in the visual cortex. Spearman’s correlation was again used to obtain the correlation scores.

All analyses described above were performed in patches of interest (POIs) taken from the BrainVoyager cortical atlas of Brodmann areas. The bilateral areas 17, 18, and 19 were used as the visual cortex POIs (the analysis presented in Fig. S2 was constrained to the left hemisphere). Analogously, the target areas in the representational connectivity analysis were also defined bilaterally. The left area 38 was used as the anterior temporal lobe POI in the analysis presented in Fig. S3. To obtain POIs that were adjusted to individual patterns of cortical folding, the participants’ cortical surface reconstructions were aligned to the atlas template in a cortex-based alignment procedure. Then, the inverse transformation matrices were used to project the atlas areas onto cortex reconstructions of each participant. The POIs were minimally dilated (one iteration of the BrainVoyager “dilate” POI function) to account for potential alignment imperfections. The analysis in POIs that were not dilated produced virtually identical results.

At the group level, the results were tested against chance levels using null distributions that were empirically derived in a permutation procedure. Specifically, each analysis was rerun 1000 times for each participant with word labels randomly assigned to this participant’s brain activity patterns in each iteration. Null distributions created in this procedure were averaged across participants and compared with the actual average results. The obtained p values (minimal value: p = 0.001) were corrected for multiple comparisons using the false discovery rate (Benjamini and Hochberg, 1995). The correction was jointly applied to all tests against chance level performed in a given analysis, in both groups. The results of correlation of neural activity patterns with the physical similarity ratings and the conceptual similarity ratings were treated as separate analyses and, consequently, corrected separately. A review of null distributions confirmed that, in all analyses, the empirically-derived chance levels were indistinguishable from a priori chance levels (i.e., accuracy = 5% in the classification analyses, Spearman’s rho = 0 in the representational similarity analyses). Thus, for simplicity, the a priori chance level is presented in the figures.

Testing for main effects and interactions were performed with ANOVAs, as implemented in SPSS 25 (IBM Corp, Armonk, NY). The Bonferroni correction, as implemented in SPSS, was used to correct the results of post hoc tests for multiple comparisons. Additional, planned comparisons testing for between-group and within-group differences were performed with two-sample and paired t tests, respectively. The Bonferroni correction was applied to the results of these tests, when appropriate, consistently with ANOVA post-hoc comparisons.

#### Searchlight analysis

The searchlight analysis was performed in surface patches of 500 vertices, using the CosmoMVPA and the Surfing toolbox (Oosterhof et al., 2011). The neural similarity matrices were created using Pearson correlation, and correlated with ratings of physical and conceptual similarity between word referents using the Spearman’s partial correlation, as was described above. The results obtained for each participant were aligned to the BrainVoyager atlas template, using forward transformation matrices obtained in the cortex-based alignment procedure. Then, the results were entered into group statistical models. One-sample t tests were used to test the results against chance level, in each group separately and when the participants from both groups were combined in one sample. Two-sample t tests and paired t tests were used to test for the differences between groups and conditions, respectively. Two-sample t tests performed on between-condition difference scores were used to test for interaction between the participant group and the condition.

CosmoMVPA was used to perform these tests. The resulting group T-maps were then converted into a threshold-free cluster enhancement (TFCE) map (Smith and Nichols 2009), calculated with standard parameters (E = 0.5, H = 2). The statistical significance of the results were determined by comparing the obtained TFCE values with null distributions obtained in the Monte Carlo simulation procedure (10 000 iterations) implemented in CosmoMVPA. The analyses were thresholded at p < 0.05, corrected for multiple comparisons (Z < 1.65). The T-values obtained in the group models were overlaid on the vertices in which the effects survived the threshold, in order to better visualize the statistical strength of the results.

#### Univariate analysis

We first performed the univariate POI analysis in the visual cortex. The analysis of contrast estimates for words presented in each modality, relative to rest periods, was performed in the same visual POIs that were used in the multi-voxel pattern analysis.

BrainVoyager was used to obtain these contrast estimates for each participant from the first-level models. At the group level, one-sample t-tests were used to test for significant effects, relative to rest. The false discovery rate was used to correct the results across all tests performed. The ANOVA, calculated in SPSS 25, was used to test for the main effects and interactions. The Bonferroni correction, as implemented in SPSS, was used to correct the results of post hoc tests for multiple comparisons. Additional, planned comparisons testing for between-group differences were performed with two-sample t tests. The Bonferroni correction was applied to the results, when appropriate.

Subsequently, the whole-brain univariate analysis was performed. The contrast estimates obtained for each participant were aligned to the BrainVoyager atlas template, using forward transformation matrices obtained in the cortex-based alignment procedure. Then, the aligned contrast estimates were entered into 2 (group) x 2 (word presentation modality) ANOVA, as implemented in BrainVoyager. All results were thresholded at p < 0.001 voxel-wise, corrected for multiple comparisons using a surface patch extent, as determined by BrainVoyager.

## Acknowledgements

This work was supported by a National Science Center Poland grant (2020/37/B/HS6/01269) and a Polish National Center for Academic Exchange fellowship (BPN/SEL/2021/1/00004) to Ł.B.

## Author contributions

M.P., M.U, and Ł.B. conceptualized and designed the study; M.P., M.U. and M.D. collected the data; M.P. and Ł.B. performed the data analyses; Ł.B. wrote the manuscript; M.P., M.U. and M.D. revised the manuscript.

## Supplementary Materials

### Supplementary Figures

**Figure S1.**
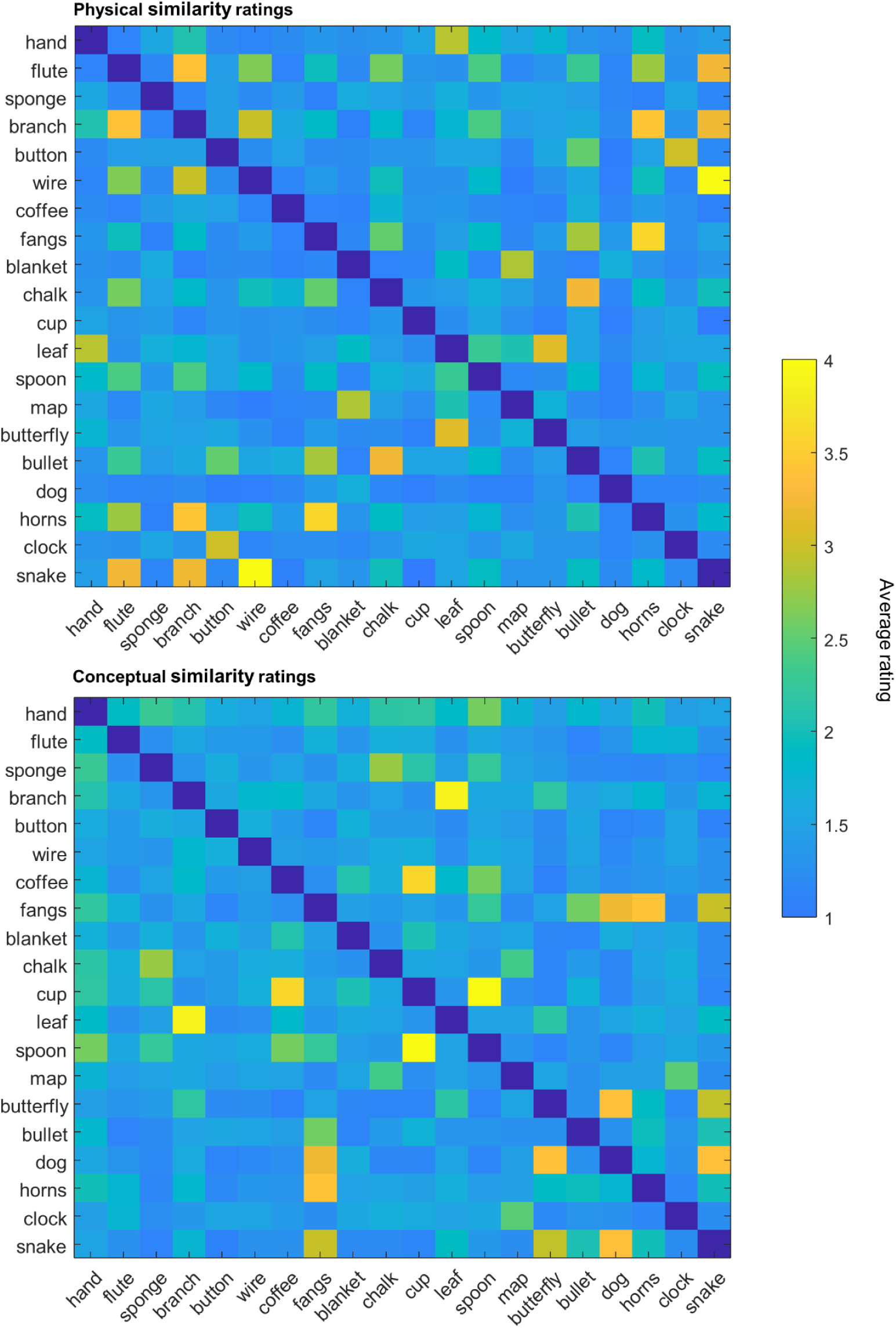
The behavioral ratings of physical and conceptual similarity between the referents of words used in the study. The matrices visualize ratings for all 190 word pairs that can be created from the 20 words used in the study. The participants rated the similarity between word referents on scales from 1 to 5. The scores were averaged across the participants (there were no average scores greater than 4). The English translations of Polish words that were actually used in the study are presented.

**Figure S2.**
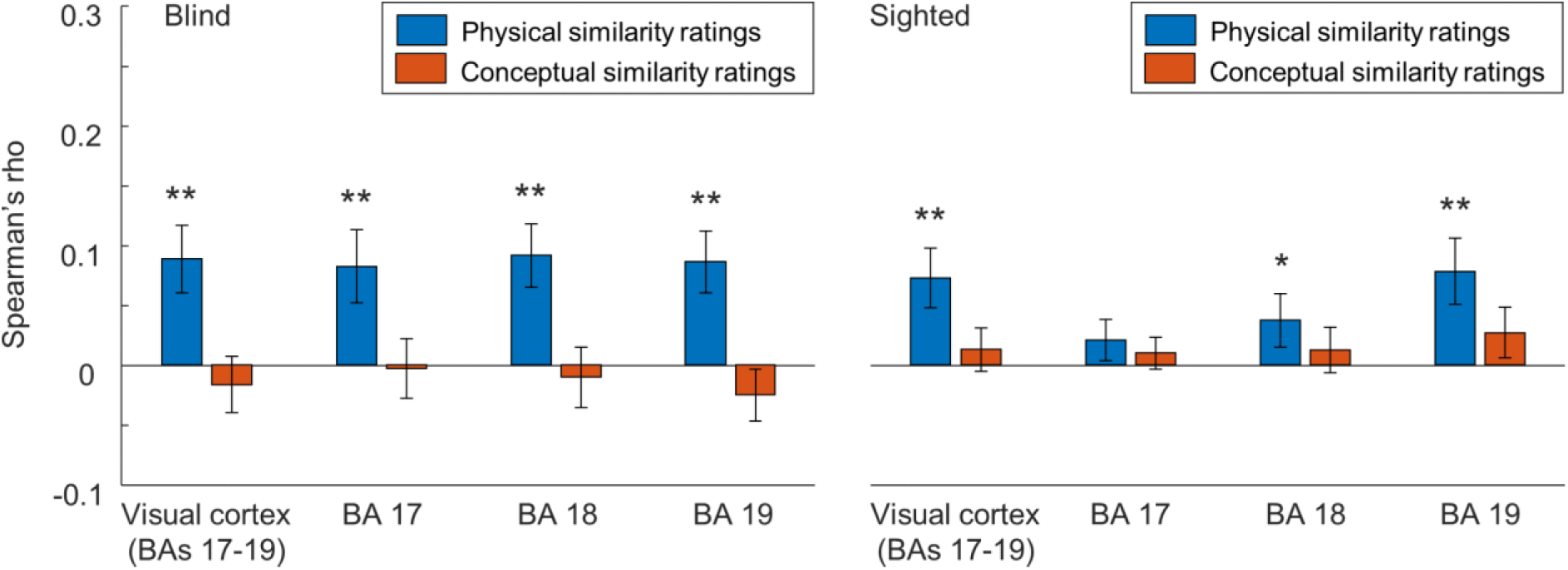
The activity patterns in the visual cortex of blind and sighted individuals reflect physical, but not conceptual similarity between word referents – analysis in the left hemisphere only. The results of correlation between the neural similarity matrices, reflecting similarity between the activity patterns for specific words in the visual areas of blind and sighted participants, and the behavioral ratings of physical and conceptual similarity between the word referents. Only activity patterns in the left hemisphere were analyzed. * p < 0.05, ** p < 0.01, corrected for multiple comparisons using the false discovery rate. Error bars represent the standard error of the mean.

**Figure S3.**
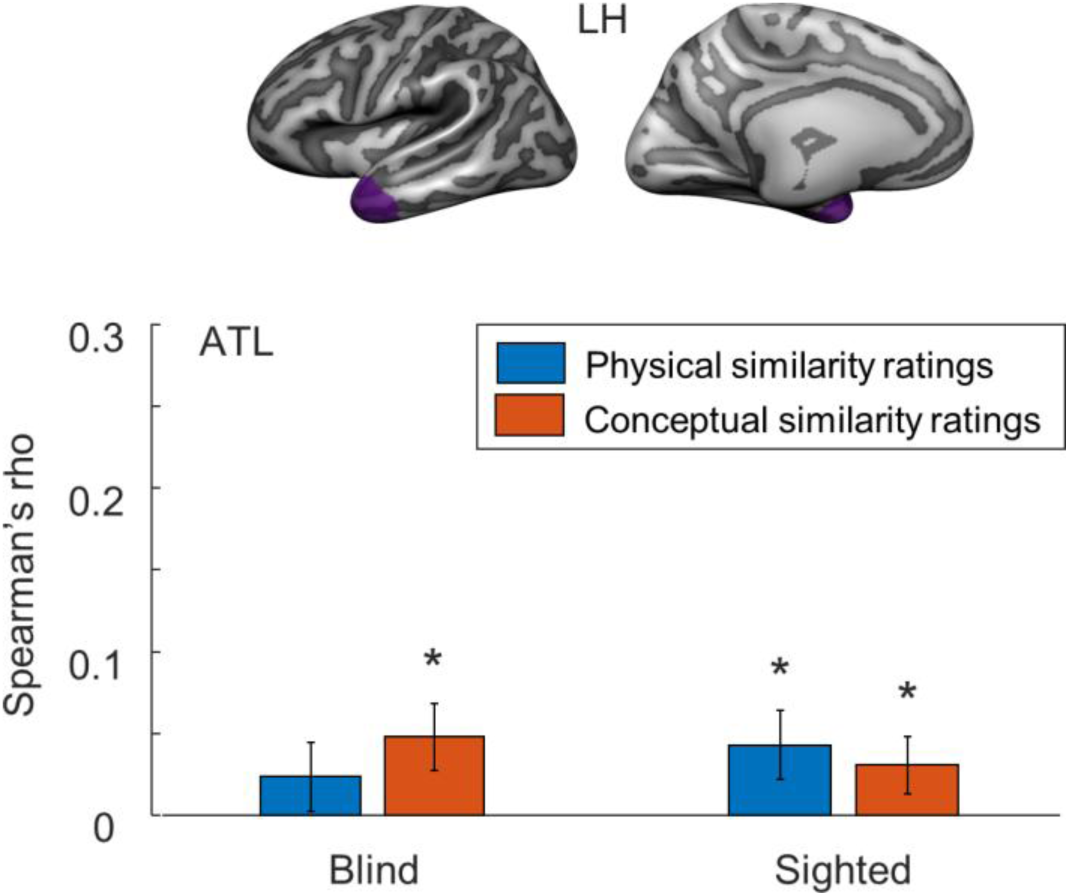
The activity patterns in the left anterior lobe reflect conceptual similarity between word referents in both participant groups. The results of correlation between the neural similarity matrices, reflecting similarity between the activity patterns for specific words in the left anterior temporal lobe (ATL, marked in purple) of blind and sighted participants, and the behavioral ratings of physical and conceptual similarity between the word referents. * p < 0.05, corrected for multiple comparisons using the false discovery rate. Error bars represent the standard error of the mean.

**Figure S4.**
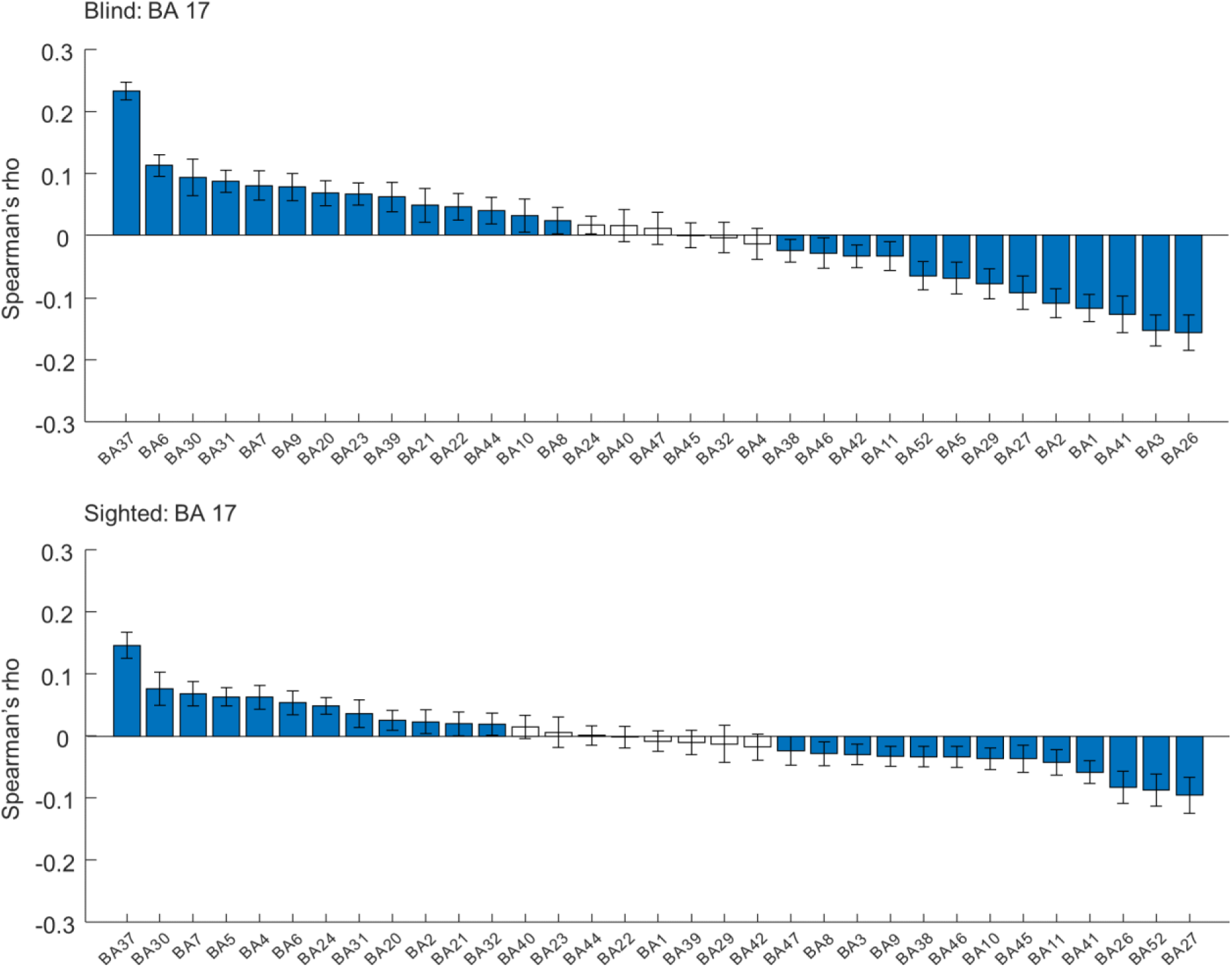
During the word processing, the visual cortex in both blind and sighted participants shows the greatest “representational connectivity” to the occipitotemporal cortex - the analysis constrained to the primary visual cortex (BA 17). For each brain region, the representational connectivity to the BA 17 was calculated by correlating neural similarity matrices, reflecting similarity between the activity patterns for specific words, obtained for the BA 17 and this region. In each participant group, the average correlation between the BA 17 and the target brain regions was subtracted from the results, to visualize regions that show higher-than-average and lower-than-average representational connectivity to the BA 17 during the word processing. The results for such regions are marked in blue (comparison against average correlation, p < 0.05, corrected for multiple comparisons using the false discovery rate). The target brain regions were sorted by the obtained representational connectivity scores. Error bars represent the standard error of the mean.

**Figure S5.**
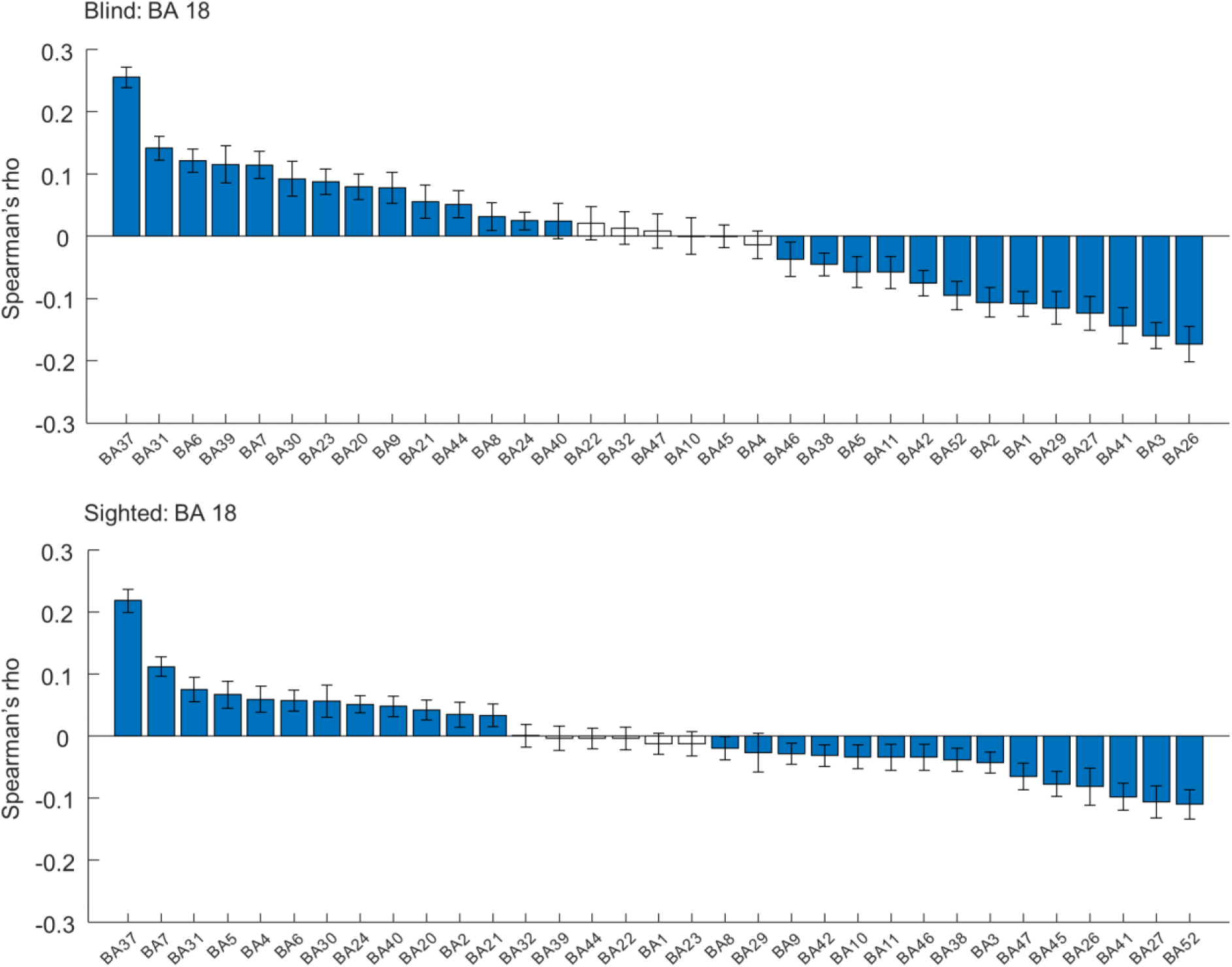
During the word processing, the visual cortex in both blind and sighted participants shows the greatest “representational connectivity” to the occipitotemporal cortex - the analysis constrained to the secondary visual cortex (BA 18). For each brain region, the representational connectivity to the BA 18 was calculated by correlating neural similarity matrices, reflecting similarity between the activity patterns for specific words, obtained for the BA 18 and this region. In each participant group, the average correlation between the BA 18 and the target brain regions was subtracted from the results, to visualize regions that show higher-than-average and lower-than-average representational connectivity to the BA 18 during the word processing. The results for such regions are marked in blue (comparison against average correlation, p < 0.05, corrected for multiple comparisons using the false discovery rate). The target brain regions were sorted by the obtained representational connectivity scores. Error bars represent the standard error of the mean.

**Figure S6.**
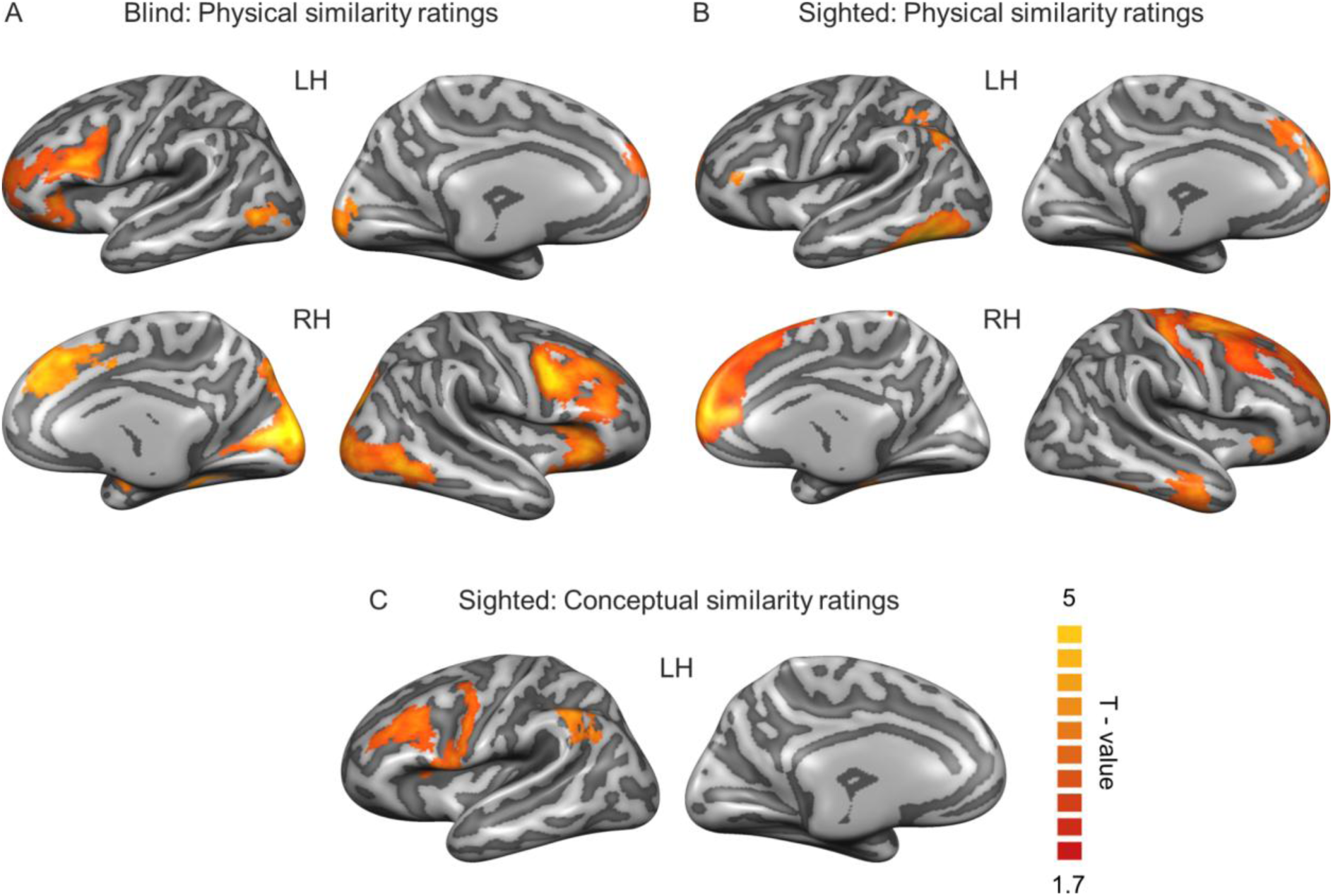
Searchlight analysis - results presented for each group separately. (A-B) The results of correlation between the neural similarity matrices, reflecting similarity between the activity patterns for specific words, and the behavioral ratings of physical similarity between the word referents (A) in the blind participants and (B) in the sighted participants. (C) Analogous analysis including the ratings of conceptual similarity between the word referents. The results for the sighted participants are presented. No significant results in the blind group were found. The statistical significance was determined using threshold-free cluster enhancement (TFCE) maps and Monte Carlo simulation. The statistical threshold was set at p < 0.05, corrected for multiple comparisons. The T-values for the effects that survived the threshold are presented for the visualization purposes.

**Figure S7.**
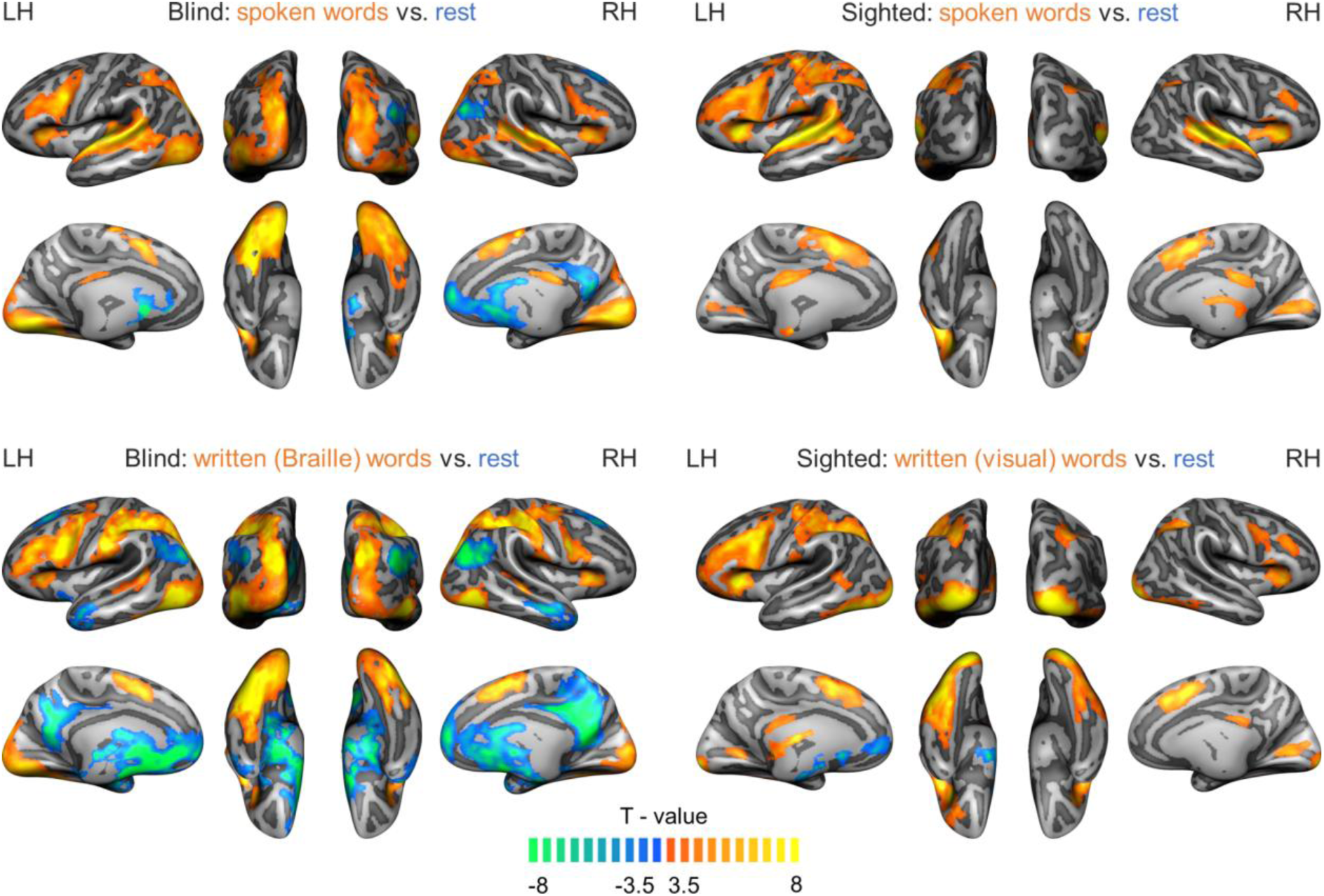
Whole-brain univariate analysis: activations for words relative to rest periods. Activations for spoken and written words, compared to rest, in the blind and the sighted group. The written words were presented in the Braille alphabet in the blind group and in the visual alphabet in the sighted group. As indicated by the font colors in the titles of the figure panels, warm colors indicate stronger activations for words, whereas cold colors indicate stronger activations for rest periods (i.e., deactivations during the processing of words). The statistical threshold was set at p < 0.001, corrected for multiple comparisons using surface patch extent.

**Figure S8.**
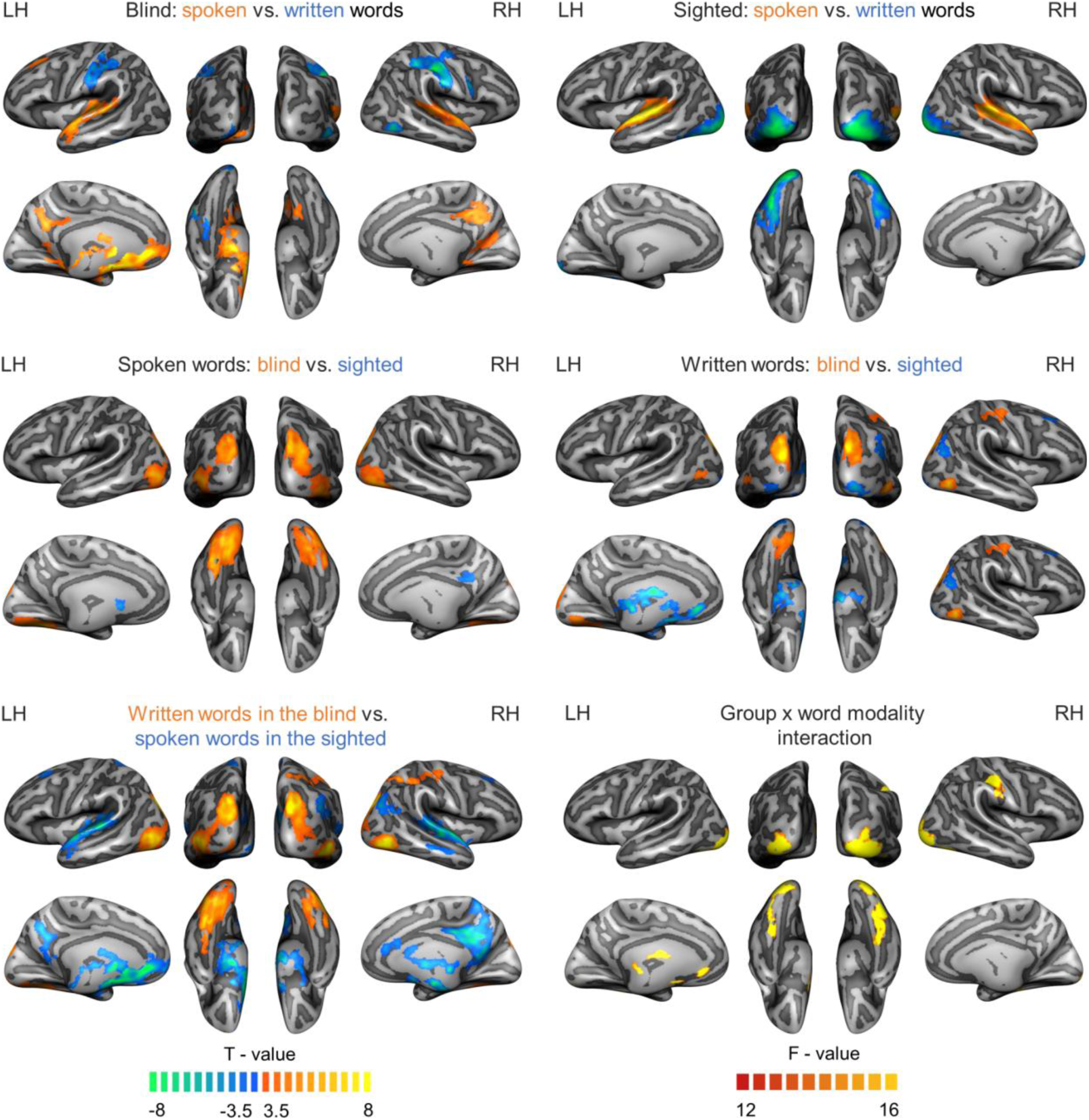
Whole-brain univariate analysis: comparisons between groups and sensory modalities of word presentations. The font colors in the titles of the figure panels match groups or conditions with warm and cold colors in the figure. The statistical threshold was set at p < 0.001, corrected for multiple comparisons using surface patch extent.

**Figure S9.**
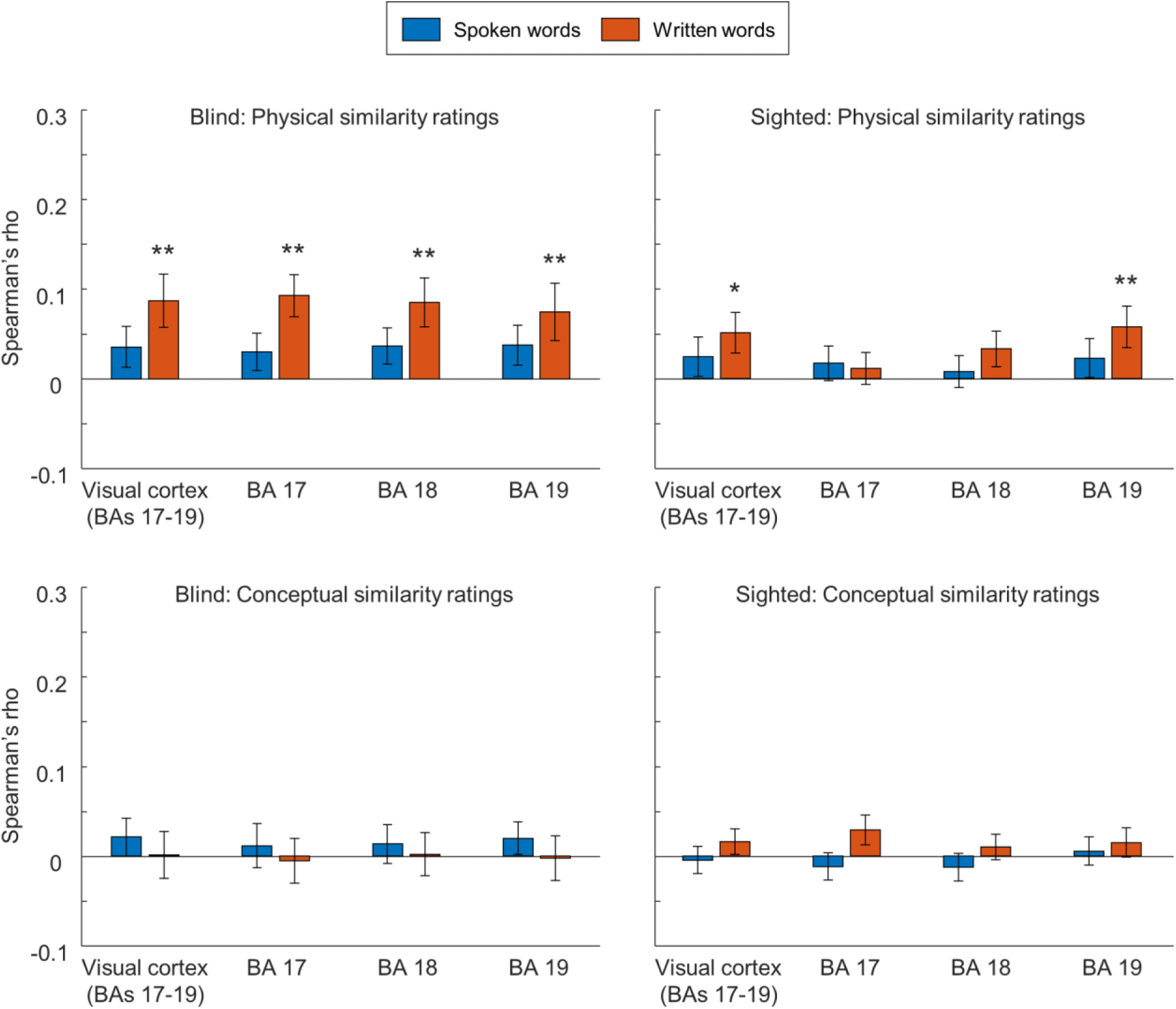
Correlation of activity patterns for words in the visual areas with behavioral similarity ratings: investigating effects of modality of word presentation. The results of correlation between the neural similarity matrices, reflecting similarity between the activity patterns for specific words in the visual areas of blind and sighted participants, and the behavioral ratings of physical (upper panels) and conceptual (bottom panels) similarity between the word referents. The analysis was run separately for runs in which words were presented in the spoken modality (blue bars) and in the written modality (orange bars). The written words were presented in the Braille alphabet in the blind group and in the visual alphabet in the sighted group. * p < 0.05, ** p < 0.01, corrected for multiple comparisons using the false discovery rate. Error bars represent the standard error of the mean.

